# First Molecular Detection of Influenza D virus in Cattle from Commercial farm in Nigeria, Sub-Saharan Africa

**DOI:** 10.1101/2023.03.15.532744

**Authors:** Abdul-Azeez A. Anjorin, Gideon O. Moronkeji, Goodness O. Temenu, Omolade A. Maiyegun, Christopher O. Fakorede, Samuel O. Ajoseh, Wasiu O. Salami, Rebecca O. Abegunrin, Kehinde O. Amisu, Kabiru O. Akinyemi

## Abstract

Influenza D virus (IDV) was first reported in pigs in the USA, and since then the virus has become a public health issue. In Nigeria, no work has been done on IDV despite the manifestation of influenza-like illness in cattle. This study aimed at molecular surveillance of IDV in cattle in Lagos. Prospective epidemiological investigation was initiated in a large commercial farm market where animals in open pens are reared, sold, and butchered under poor hygienic conditions without adequate biosecurity measures. A total of 80 nasopharyngeal swabs were collected between October and November 2021. The samples were extracted using an RNA purification kit (NIMR). RNA extracts were amplified following a two-step PCR using FIREScript RT cDNA synthesis kit (Solis Biodyne, Estonia), followed by PCR OneTaq Quick-load 2x master-mix (NEB, UK) in a Rotor-Gene thermocycler (Qiagen, Germany). Amplicons were detected using a 1.5% Gel electrophoresis. IDV was detected in 26/80 (32.5%) cattle. Sick animals showed 65% (17/26) almost double burden of IDV higher than 34.6% (9/26) in a healthy population, including 88.2% (15/17) Cattle with diarrhoea and 11.8% (2/17) with nausea having IDV. Bull recorded more than twice, 18/26 (69.2%) incidence by gender compared to Cow. Age prevalence showed 62.23% (18/26) highest detection in cattle of 4 years old, followed by 23.07% (6/26) in 5 years old, while the lowest 7.69% (2/26) was recorded in 3 years old. This study showed the first molecular detection of IDV in Nigeria and West Africa sub-region to the best of our knowledge. It underscores the need for continuous surveillance of IDV at the animal-human interface.

## INTRODUCTION

The intricacies of bovine respiratory disease (BRD) are alarming to public health and have consequential devastating economic losses, especially on young bovines, livestock weight reduction, dairy products, and overall farm profitability globally (DeDonder and Apley, 2015, Pastrana et al., 2022). Different viruses have been incriminated in BRD, including bovine alphaherpesvirus 1, bovine coronavirus, bovine viral diarrhoea virus, bovine parainfluenza type 3 virus, and bovine respiratory syncytial virus (Salem et al., 2019, Pastrana et al., 2022).

A new influenza D virus (IDV) identified to infect cattle was added to the BRD-associated viruses (Nissly *et al*., 2020). IDV was first isolated in April 2011 in Oklahoma, USA from a frazzling swine and thereafter in cattle with respiratory disease (Hause et al., 2013, Hause et al., 2014). This virus isolated from pigs caused influenza-like symptoms, with up to 50% overall homology to the human influenza C virus (Hause et al., 2013). It was found to be an envelope, single-stranded, negative-sense RNA virus of the family Orthomyxoviridae. It has seven genomic segments encoding 9 proteins (NS1 and NEP) (Hause et al., 2013, Hause et al., 2014, Collin et al., 2015).

According to the International Committee on Taxonomy of Viruses (ICTV) 2021 ratification, the influenza virus now belongs to the new Realm- Riboviria, Kingdom- Orthornavirae, Phylum- Negarnaviricota, Sub-phylum- Polyploviricotina, Order- Articulavirales, Family-Orthomyxoviridae, Class- Insthoviricetes, Genus- Alpha, Beta, Delta, and Gamma (formally C) influenzavirus, and Species- Influenza A, B C, and D virus. However, based on the hemagglutinin-esterase (HEF) gene, there are 3 main antigenic and genetic clusters in circulation to date: D/OK which is known worldwide; D/660 which is only in the USA and recently confirmed in Europe, and D/Japan currently in Japan (Odagiri et al., 2018).

There had been successful detection of IDV in Asia, Europe, Central and North America, and some parts of Africa (Hause et al., 2013, Jiang et al., 2014, Collin et al., 2015, Chiapponi et al., 2016, Murakami et al., 2016, Sanogo et al., 2021). Also, IDV has been reported in other hosts like buffalo, camel, ferret, goat, and sheep in Togo, Kenya, and China (Salem et al., 2017, Zhai et al., 2017) and small ruminants, equine and feral swine (Quast et al., 2015, Ferguson et al., 2015, Nedland et al., 2018). Low seroprevalence of less than 2% has been reported in humans, but this may be due to previous exposure to the influenza C virus that shares the same surface antigen, haemagglutinin esterase fusion (HEF) with IDV, a distinguishing structural feature from influenza A and B subtypes that possess 2 surface glycoproteins-haemagglutinin and neuraminidase for viral attachment and release respectively.

IDV transmission occurs either through direct or indirect spread via aerosol whereby infected animals shed viral particles in nasal secretions through airborne to other animals. Primarily, IDV becomes garner from close contact with an infected animal (Hause et al., 2013). Replication does occur within the upper respiratory tract with no trace in the trachea and lungs (Hause et al., 2014). Clinical manifestations usually occur after the incubation period of 1-4 or more days (Hause et al., 2013). Isolation of the virus is the main direct diagnosis of IDV (Ferguson et al., 2016) and PCR remains the most commonly used method due to its specificity and sensitivity and the duration of time to complete the assay (Faccini et al., 2017).

Continuous surveillance and molecular techniques have increased the global epidemiology of IDV. Huang et al. (2021) revealed a molecular prevalence of 10% from 100 nasal swab samples screened in cattle in California, USA between 2018 and 2019. Another study from the University of Calgary, Veterinary Sciences reported a molecular prevalence rate of 22.8% from 232 cattle (Zhang et al., 2019). In the UK, a molecular prevalence rate of 8.7% was reported from 104 cattle nasal swab samples screened for the presence of IDV (Dane et al., 2019). In Asia, 12.8% IDV was detected from 156 dairy cattle and 7.3% from 55 native yellow cattle in China (Zhai et al., 2017). Limited studies in Africa revealed 3.07% incidence rate of IDV from 2 positive nasal swab samples out of 65 cattle tested in Namibia (Molini et al., 2022).

Therefore, to ameliorate the possible challenge of epizootic with the potential zoonotic transmission based on the 1.3% seroprevalence of IDV in the general human population (Hause et al., 2013), there is a need for continuous surveillance of IDV for proper documentation that will inform the government including the policy-makers on the need for more preparedness at the animal or reservoir host interface, especially in a country like Nigeria with over 20 million cattle per head that are reared indiscriminately in the backyard and commercial farms. In essence, this study was aimed at investigating IDV in cattle on the largest commercial farm in the most populous city in Africa, Lagos State, Nigeria.

## MATERIALS AND METHODS

### Study design and site

A commercial farming-based molecular study and prospective epidemiological surveillance were carried out on cattle in the Kara market between October and November 2021. Kara cattle market is an international market with commercial activities of notable ruminant animals of over 20,000 per week. It houses an abattoir for slaughtering and sales of raw beef daily. It is located along the Lagos-Ibadan expressway on the return journey to Lagos in Berger. The settlement is situated beside the flow-through of the Ogun River (Fig 1) with a bridge linking Lagos and Ogun States.

**Fig 1:**
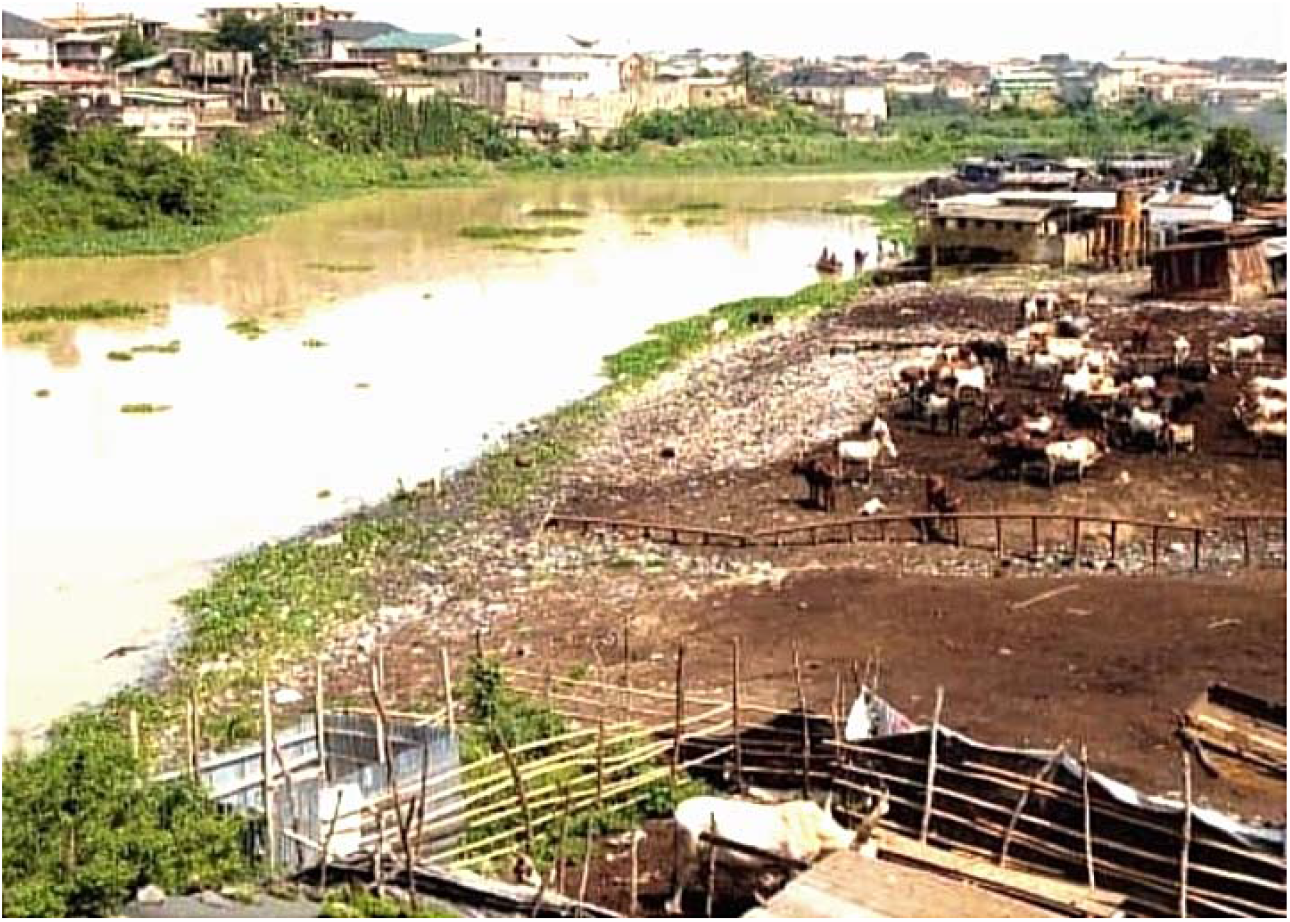
Kara market flow of Ogun River. https://maps.app.goo.gl/3yRMQwjiCxk4Q8XL8.

The cattle colony has been in existence for over 34 years with mixed ethnic groups of Fulani, Hausa, and Yoruba tribes of about 700 people trading cattle and other livestock like sheep and goats of different species. The ruminant animals irrespective of the species are usually sourced and transported from the northern part of Nigeria weekly in heavy-duty long trailers along with the herders without any biosecurity measures.

The market lacks good hygiene practices with the floor covered with dung(Fig 2a) of cohabiting animals that are usually discharged into the water body of a river flow-through that also serves as the major source of water for the farm (Fig 1).

**Figure 2a.**
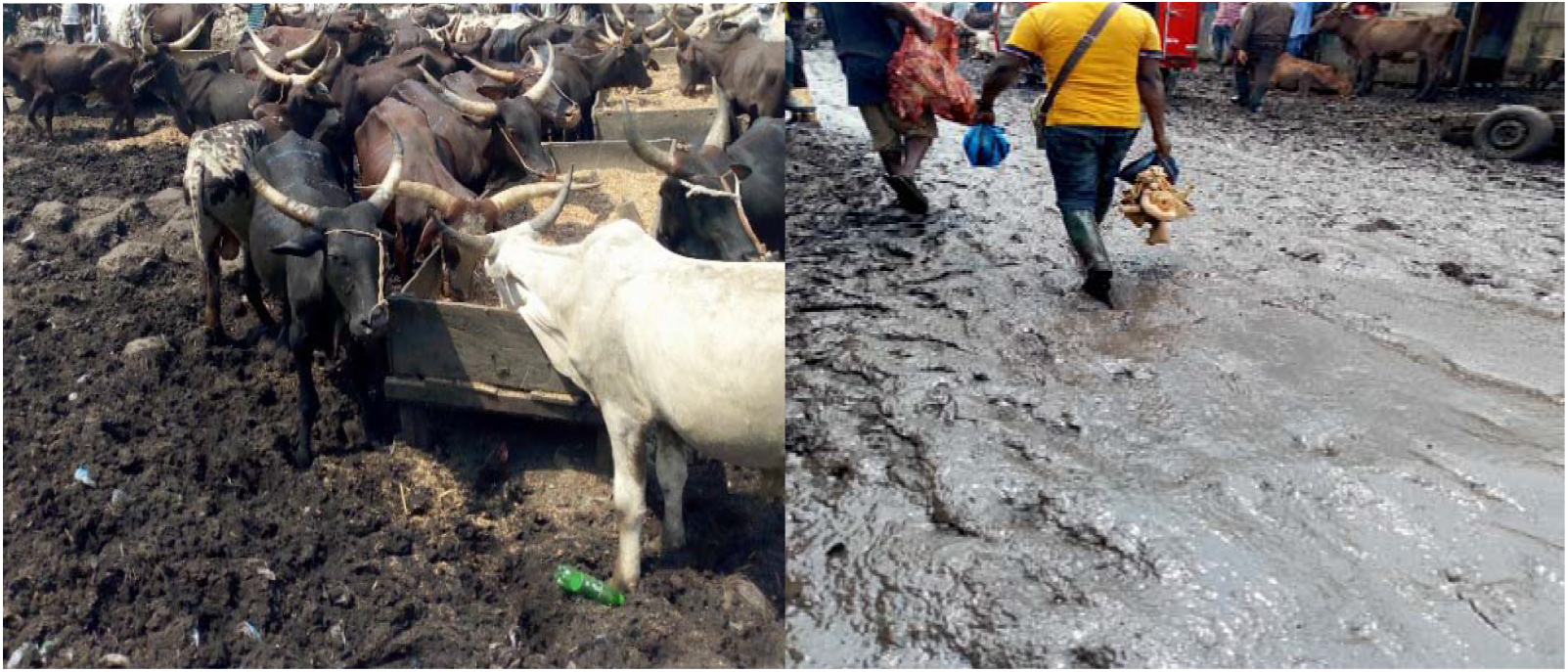
Cattle dung covers the ground as the carpet at the cattle feedlot during the dry season at Kara market. **b**. Marshy ground during the rainy season at Kara market.

The market is normally marshy during the rainy season with resultant difficulty in the movement for both herders and buyers (Fig 2b).

### Inclusion and exclusion criteria/ethical approval

Kara cattle market was selected for this study considering its large cattle population as the biggest commercial farming center in southwestern Nigeria and based on the farm’s proximity to the molecular research laboratory. Sick cattle were particularly targeted and recruited from their herders to increase the chances of detecting viral RNA along with healthy animals for possible comparison. Farm herders who were reluctant or those who refused their cattle to be sampled due to fear of the unknown despite explanations by Veterinary doctors were excluded.

Ethical approval was obtained from the Lagos State University Teaching Hospital (LASUTH) Research and Ethics Committee with the registration number NHREC04/04/2008, and reference number LREC/06/10/1272 respectively.

### Sample collection and RNA extraction

By engaging different herdsmen, purposive sampling was carried out on various cattle feedlots. A total of 80 samples were collected with dragon swab sticks having flexible plastic shafts. The swab was inserted gently and horizontally into the nostril to the nasal cartilage while it was rotated laterally to absorb secretion. Each swab sample was transferred into a pre-labeled cryovial tube containing 1ml of normal saline as a viral transport medium (VTM).

The cryovials were transported in batches in a cold chain to influenza and other respiratory tract viruses (IORTV) research facility in the molecular research laboratory, Department of Microbiology, Lagos State University and were kept in a 4°C refrigerator before extraction time. Aliquots of all samples were frozen at -30°C.

Four samples were pooled into one tube to give a total of 20 pooled samples for an effective cost of reagents, laboratory analysis, and time management. Viral RNA was extracted using an RNA extraction kit (NIMR, Nigeria). All positive pooled samples were re-extracted as separate samples. Two-step PCR was done, with an initial cDNA conversion using Solis Biodyne (FIREScript RT cDNA Synthesis KIT, Estonia). This was followed by PCR using One Taq Quick-Load 2x Master Mix with Standard Buffer from BioLabs (UK).

### Master mixture preparation and PCR amplification

The master mix was prepared for cDNA from the RNA extracts following the manufacturer’s instructions. Briefly, for a single reaction, 0.5µl dNTP MIX, 2 µl of 10x RT reaction buffer with DDT, l0.5 µl of RiboGrip RNase Inhibitor, 20 µl of Nuclease-free Water and 1 µl of FIREScript RT were dispensed each into RNase free 0.2ml PCR strip tube, which was followed by addition of 1µl each of oligo (dT) and random primers for cDNA synthesis. Thereafter, 5µl of extracted genome template of the sample was added. Thermocycling conditions were carried out in a Rotor-Gene Q PCR machine (Qiagen, Germany) with the following optimization: primer annealing at 25°C for 600 secs, Reverse transcription (RT) for 55°C for 1200 secs, with enzyme inactivation at 85°C for 300 secs.

Furthermore, a second step of PCR amplification was performed on the cDNAs. According to Chiapponi et al. (2019), IDV-specific primers- IDVF: 5- TGG ATG GAG AGT GCT GCT TC -3 and IDVR 5- GCC AAT GCT TCC TCC CTG TA -3 were synthesized from STAB Vida, Portugal. For a single reaction, 12.5 µl of One Taq Quick-load 2x master mix with standard buffer (New England BioLabs, UK) was aliquoted into a DNase-free 0.2ml transparent PCR tube. Forward and reverse primers of 0.5 µl each was added to the aliquot, followed by 6.5 µl of nuclease-free water to make-up a 20 µl each per reaction. The master mix was short-spun by vortexing before 5µl of cDNA was appropriately added to each tube prepared. The thermocycling condition included: initial denaturation at 94°C for 30 secs, 30 cycles at 68°C for 60 secs, and final extension at 68°C for 300 secs.

### Amplicon detection by gel electrophoresis

PCR amplicons were detected by gel electrophoresis. Agarose gel electrophoresis (1.5%) was prepared by measuring 3g agarose powder that was dissolved in 200 ml of 50x TAE buffer in a uniform solution. The solution was microwaved for about 3-5 mins. It was allowed to cool uniformly by continuous swirling to about 40-50°C before the addition of 3ul ethidium bromide stain. The agarose was poured into a casting tray with an inserted comb to create the required number of wells. The comb was removed while the wells were loaded with premixed 5ul of sample amplicons and 1ul of 6x loading dye. A total of 3ul of a 100bp DNA ladder was added for each run. The agarose gel was run at 130V and 400 amperes for 40mins using a gel electrophoresis machine (Chemikalien Laborbedarf, Germany). A transilluminator imager (Dark Reader transilluminator, USA) was used to view and capture the agarose gel images for proper documentation, in a dark room. All the laboratory procedures were carried out following the manufacturer’s instructions.

### Statistical analyses

Epidemiological data were computed into contingency tables and analyzed with Chi-square (and Fisher’s exact) test using GraphPad Prism 9 for Windows Version 9.0.0 (121) (GraphPad Software Inc., San Diego, CA, USA). P-values were measured and statistically significant differences were determined for each epidemiological parameter including age, gender, and health status of healthy and sick animals presenting with clinical symptoms collected with a paper-administered questionnaire to the herders. No adjustments were made for missing data. P-values were two-sided, and a significant difference was considered at P value <0.05.

## RESULTS

Out of the 80 nasal swab samples tested by PCR, 26 samples were positive (Figures 3-4), resulting in a prevalence rate of one in three cattle (32.5%) having influenza D virus (Figure 5). The molecular prevalence data by health status showed that the burden of IDV almost doubled by 17/26 (65%) in sick cattle in comparison to 9/26 (34.6%) in the healthy group. Out of the sick cattle, 15/17 (88.2%) that recorded diarrhoea and 2/17 (11.8%) having nausea were positive for IDV (Table 1). Based on gender distribution, the incidence of IDV was more than twice higher, 18/26 (69.2%) in Bulls compared to 8/26 (30.7%) in Cow (Table 2). The distribution of IDV prevalence by age of cattle showed the highest result in 4 years old cattle 18/26 (62.23%), followed by 5 years 6/26 (23.07%) and 3 years with the lowest rate of 7.69% (2/26) (Table 3).

**Figure 3:**
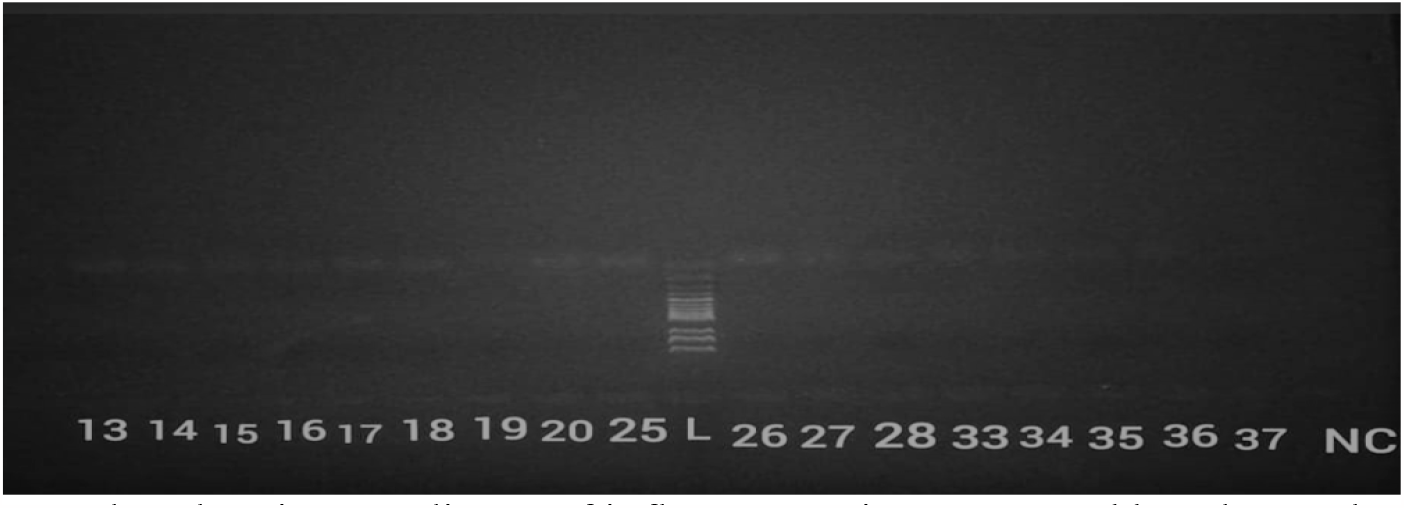
Panel A showing amplicons of influenza D virus separated by Electrophoresis with positives on lanes: 13, 14, 15, 16, 17, 18, 20, 25, 26, 27, 28, 33, 34, 35 and 36, as against NC-negative control.

**Figure 4:**
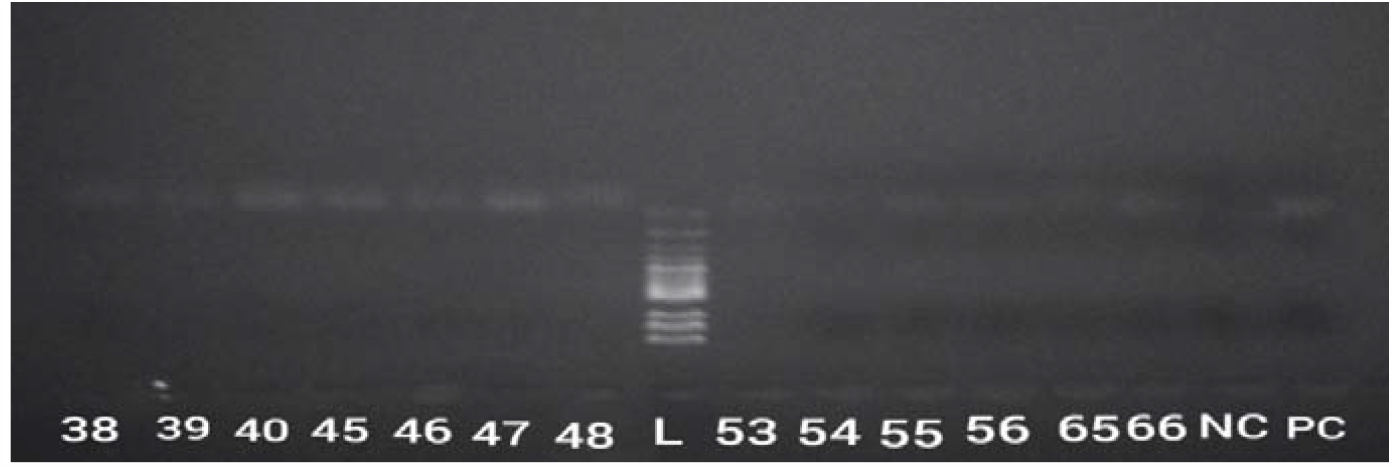
Panel B depicts positive amplicons of the Influenza D virus on lanes: 38, 39, 40, 45, 46, 47, 48, 55, 56, 65, and 66. NC- negative control while PC- positive control.

**Table 1:** Molecular prevalence of influenza D virus in cattle based on health status in Lagos

| Health status | Total N= 80 (%) | Positive n=26 (%) | P value |
| --- | --- | --- | --- |
| Sick | 39 (48.8) | 17 (65.0) | <0.0001 |
| Healthy | 41 (51.3) | 9 (34.6) |  |
|  | **N= 26** | **n=17** |  |
| Diarrhoea | 15 (57.7) | 15 (88.2)* |  |
| Nausea | 2 (7.7) | 2 (11.8)* |  |

\*Diarrhoea and nausea percentages were derived from the total sick positive denominator of 17.

**Table 2:** Incidence of influenza D virus in cattle based on gender in Lagos, Nigeria

| Gender | Total N=80 (%) | Positive n=26 (%) | P value |
| --- | --- | --- | --- |
| Bull | 56 (70) | 18 (69.2) | >0.9999 |
| Cow | 24 (30) | 8 (30.7) |  |

**Table 3:** Distribution of influenza D virus in cattle based on age in Lagos, Nigeria

| Age (yrs.) | Total N= 80 (%) | IDV Positive (%) | P value |
| --- | --- | --- | --- |
| 3 | 3 (3.8) | 2 (7.7) | 0.11 |
| 4 | 47 (58.8) | 18 (62.2) |  |
| 5 | 30 (37.5) | 6 (23.1) |  |

## DISCUSSION

This study accounted for the first molecular prevalence of influenza D virus in Nigerian cattle among both healthy and sick populations. The result showed a high 34.6% rate of IDV even among healthy cattle with no symptoms of IDV infection. It was discovered that the circulation of IDV in cattle cohorts widely spread among commercial cattle that are ready for butchering in the abattoir. The incidence rate among sick cattle at 65% was about two times compared to the 34.6% molecular prevalence detected in healthy IDV carriers.

A previous study done in the Southern African country of Namibia by Molini et al. (2022) showed a molecular prevalence of 3.1% from 65 cattle which is far low when compared to the current study. This study was also higher than the prevalence of 22.8% reported elsewhere in Canada by Zhang et al. (2019); 19% in China by Zhai et al. (2017); 10% in the USA by Huang et al. (2021), 8.7% reported in the UK by Dane et al. (2019), 5.6% in Ireland by Flynn et al. (2018), 4.5% reported from France by Ducatez et al. (2015), 3.9% in Turkey by Yilmaz et al. (2020) and the lowest 1.3% documented in Italy by Chiapponi et al. (2019).

The highest general molecular prevalence of 32.5% of IDV reported in Cattle in Africa from our study calls for concern. We, therefore, looked closer at the farm settlement to probe likely predisposing factors. A myriad of favourable conditions enhancing viral transmission between the cattle herds was discovered on the farm. Other ruminant animals were sold in the same market location, along with critical promoters of viral spread including unhygienic environmental factors, lack of biosecurity and biosafety measures, decrease in proportional diet that could boost the immune response against viral invasion, the influx of different species of ruminant animals that can harbour influenza virus, and inadequate health management for the herds. This agrees with the relationship of BRD with the host, pathogen, animal management, and environmental factor earlier posited by Chiapponi et al. (2019).

Efforts must therefore be put in place to reverse the above-mentioned trends in all herd farms and markets in other to curb the proliferation of IDV and other bovine viruses. Furthermore, deliberate attempts must be made to avert transmission of IDV to humans through occupational hazards due to the lack of biosecurity measures as with other influenza subtypes especially influenza A that caused the swine influenza pandemic of the 21^st^ century. This is important as previous studies by Hause et al. (2013), Hause et al. (2014), Quast et al. (2015) have reported successful replication and isolation of IDV in the lower and upper respiratory tracts of other mammals, along with the IDV cell receptor protein binding structure similarity having 50% homology to human influenza C virus. Consequently, further studies on public health threats of IDV especially in one health are critical.

The burden of IDV reported in sick animals, almost doubling that of the healthy population may be accounted for by viral shedding in the nasopharynx of the sick animals, as sick animals have been found to readily shed viruses before the manifestation of clinical signs and symptoms in the host (Anjorin et al., 2017). Other studies have also reported IDV-positives from both healthy cattle and animals manifesting clinical pathologies (Flynn et al., 2018). Hence, sick animals may pose danger to the farm with a continuous circulation of the virus to new herds and even farm workers. White et al. (2016) reported a very high 86% IDV antibodies in farm workers after exposure by working with sick cattle. The high percentage of Cattle that came down with diarrhoea followed by nausea call for serious attention as these normally result in loss of fluids and off-feed, leading to weight loss common to BRD, stunted growth, and consequential profit lost by the farmers. This also affects the industrial sector, notably the dairy industry. We strongly recommend the isolation of such sick animals once they are suspected.

The gender disparity observed with the Bulls recording more than twice the incidence compared to Cows may be due to different reasons including sterility factor usually performed for taste enhancement that often alters hormonal secretions in the male animals. This aligns with a previous report of the higher molecular prevalence of the influenza virus in other female mammals including pigs by Anjorin et al. (2017). Age prevalence study showed the highest distribution of IDV in more than 60% of cattle in the middle-aged animals with the lowest recorded in the youngest. This can be so as the middle-aged animals are expected to have lost their protective maternal antibody unlike the younger-aged animals and yet, have immature immunity compared to the oldest group.

This study is however without its limitations, including the limited number of animals sampled on a single, though a major farm in the region, and did not take into account circulation of IDV based on seasonality. Hence, in West Africa and Nigeria in particular, further research is highly warranted for continuous molecular surveillance of one health including the need for genomic sequencing of the influenza D virus in cattle to avert the potential danger of future antigenic and genomic changes.

## CONCLUSION

This study documented for the first time, molecular detection of influenza D virus (IDV) in Cattle in Lagos, Nigeria, and West Africa, to the best of our knowledge. A general incidence of 32.5% was reported with a high IDV prevalence of 34.6% among healthy cattle. Observation from this study highlights the importance of IDV as an agent of respiratory tract infection in cattle in Lagos. In-depth studies are required to exactly assess the economic impacts of IDV infections on commercial livestock markets and the occupational risks among farm workers.

## Acknowledgments

The abstract for this study was presented at the 7th World One Health Congress (WOHC) Conference held at the Sands Expo & Convention Centre, Singapore from 7 – 11 November 2022 under abstract number 15241. The presentation of this work at the WOHC/Singapore was supported with CEPI global south travel grant, 2022 awarded to AAA.

We hereby acknowledge Alexander Von Humboldt (AvH), Germany for the equipment grant support awarded to KOA, which was used in the procurement of a Rotor-Gene thermocycler, cold-chain centrifuge, Gel-documentation system, and other accessories used in this study at the Department of Microbiology Molecular biology research laboratory, Lagos State University.

## Author Contributions

**Abdul-Azeez A. Anjorin**: Performed study design, carried out laboratory and statistical analysis, performed literature search, wrote and edited the manuscript, and supervised the work.

**Gideon O. Moronkeji, Goodness O. Temenu, Omolade A. Maiyegun**: Carried out sample collection, reagent procurements, literature search, partook in laboratory analysis, and participated in the manuscript draft.

**Christopher O. Fakorede, Samuel O. Ajoseh, Wasiu Salami, Rebecca O. Abegunrin**: Instrumentation, and laboratory technical support, performed literature search, edited the manuscript and did reference sorting.

**Kehinde O. Amisu, Kabiru O. Akinyemi**: Equipment procurement, laboratory technical guide, edited the manuscript, and gave general support.

## Data Availability Statement

All relevant data have been included in the manuscript.

